# Developmental stage-dependent effects of perceived predation risk on physiology and fledging success of tree swallows (*Tachycineta bicolor*)

**DOI:** 10.1101/2022.12.27.522041

**Authors:** Sabrina M. McNew, Conor C. Taff, Cedric Zimmer, Jennifer J. Uehling, Thomas A. Ryan, David Chang van Oordt, Jennifer L. Houtz, Allison S. Injaian, Maren N. Vitousek

## Abstract

The risk of predation directly affects physiology, behavior, and fitness of wild birds. Social interactions with conspecifics may affect how individuals respond to stressors such as predators. Strong social connections could help individuals recover from a stressful experience; however, competitive interactions also have the potential to exacerbate stress. Few studies have investigated the interaction between environmental stressors and the social landscape in wild bird populations. Here, we experimentally simulated predation attempts on breeding female tree swallows (*Tachicyneta bicolor*). At the same time, we manipulated female breast plumage color, a key social signal. Simulated predation events on tree swallows negatively affected their nestlings’ condition, telomere lengths, and fledging success. However, the effects of experimental manipulations were timing-dependent: simulated predation during the early nestling period was more detrimental than “predation” during incubation. Contrary to our expectations, manipulation of the social environment did not affect the response of tree swallows to simulated predation. However, manipulating female plumage during the nestling period did affect nestling size, indicating an effect of the social environment on reproductive success. Our data demonstrate that transient stressors on breeding female birds can have carry-over effects on their nestlings, some of which may be long-lasting.

## INTRODUCTION

Predation is an important source of mortality for birds (Martin, 1993; Ricklefs, 1989). Nestlings are highly vulnerable while in the nest and behaviors such as incubation and provisioning also expose parents to predation. Even in the absence of direct consumption, the perceived risk of predation impacts behavior, physiology, and fitness of adults and their nestlings (Lima, 2009; Zanette et al., 2011). Predation therefore is not just an acute challenge that forces temporary changes in physiology and behavior; it also creates a “landscape of fear” that may affect individuals’ life-history and even population dynamics (Brown et al., 1999; Clinchy et al., 2013; Laundre et al., 2010).

Animals vary in their resilience to predation, i.e., their ability to withstand this stressor and return to normal functioning following a confrontation with a predator (Davis et al., 2021). One potential factor influencing resilience is social connectivity. For example, baboons (*Papio hamadryas ursinus*) respond to the loss of a close family member to predation by increasing social grooming, which may help lower stress hormone levels back to baseline (Engh et al., 2006). Social position and connectedness have emerged as key mediators of the psychophysiological effects of stress (Charuvastra and Cloitre, 2008; Holt-Lunstad et al., 2010; Yang et al., 2016). However, the cognitive and behavioral costs of non-consumptive predation are difficult to observe and measure in natural populations (Clinchy et al., 2013) and studies of the relationship between social connectivity and stress resilience have been conducted primarily in primates (Creel et al., 2013).

The physiological mechanisms involved in the stress response are key to understanding stress resilience and the long-term consequences of stressors. A major component of the stress response is the hypothalamic—pituitary—adrenal (HPA) axis, which regulates glucocorticoid hormones (Sapolsky et al., 2000; Wingfield et al., 1998). The release of glucocorticoid hormones helps an organism maintain fitness by mobilizing energy stores and shifting resources away from reproduction, growth and maintenance and towards an “emergency” behavioral and physiological state focused on surviving the immediate threat (Wingfield et al., 1998). Although this physiological stress response is an important adaptation allowing animals to react rapidly to challenges, chronic exposure to glucocorticoids can have negative fitness consequences (Sapolsky et al., 2000).

Glucocorticoids may connect stressors to fitness through effects on telomeres (Haussmann and Heidinger, 2015; Haussmann and Marchetto, 2010). Telomeres are repetitive sections of non-coding DNA that “cap” the ends of chromosomes and help maintain chromosome integrity during replication. Telomeres degrade over the course of an animal’s lifetime and telomere shortening is associated with disease and senescence (Angelier et al., 2018; Asghar et al., 2015). Both genetic and environmental factors influence telomere length and so telomeres are used as a proxy for “long-term somatic state,” an integrative measure of an individual’s condition (Benowitz-Fredericks et al., 2022). Chronically high levels of glucocorticoids increase somatic damage from inflammation and oxidative stress, which are also linked to telomere loss (Angelier et al., 2018; Ridout et al., 2018). Early life stress may have particularly strong effects on telomeres (Injaian et al., 2019; Ridout et al., 2018; van Lieshout et al., 2021). For instance, nestling European shags (*Phalacrocorax aristotelis*) exposed to simulated predation events experienced higher stress-induced corticosterone (the main glucocorticoid in birds) concentrations and increased telomere loss over the course of the experiment (Herborn et al., 2014). Juvenile telomere lengths can predict overall lifespan (Haussmann et al., 2005), thus even transient stressors early in life may shorten overall life expectancy.

In this study we exposed breeding female tree swallows (*Tachycineta bicolor*) to two different experimental treatments: simulated predation and manipulation of the social environment. Heightened predation risk is associated with changes in parental care of tree swallows (Wheelwright and Dorsey, 1991) and changes in investment of within-pair vs. extra-pair young (Hallinger et al., 2020). Previous experiments that manipulated perceived predation risk in our population showed that breeding females that had a robust glucocorticoid response along with a strong negative feedback to predator exposure were less likely to abandon nests during incubation (Zimmer et al., 2019). However, it is not clear whether the effects of predation carry over to nestling condition or physiology. If predation stress on breeding females is transmitted to her offspring, it could affect the nestlings’ telomere lengths and overall lifespan.

In addition to simulating predation, we manipulated the social environment by dulling the white breast plumage of females. In tree swallows, breast plumage is an important signal and naturally brighter plumage is associated with greater immunity, reproductive success, and the frequency of social interactions at the nest (Beck et al., 2015; Taff et al., 2019a). A previous experiment in our population found that manipulating female breast brightness – in the absence of a separate environmental stressor – changed the patterns of social interactions and led to higher reproductive success for dulled females, particularly when those females were initially bright (Taff et al., 2019b). Thus, signal variation alone creates feedback between the social environment, physiology, and fitness and may help mediate resilience to external stressors.

We tested for the effects of simulated predation and social manipulation on nestling growth, physiology, and fledging success during one field season (2018). The following year (2019), we repeated the experiment, switching the order of the treatments between years to test how timing of stressors affected nestling outcomes (Fig. 1). We predicted that elevated predation stress on mothers would lead to reduced parental care and overall poorer nestling outcomes. Second, we expected that there would be a tradeoff between the stress response and telomeres: nestlings that responded to predation stress with an elevated stress response would have shorter telomeres. Finally, we expected that experimentally dulled females would be less resilient to environmental stressors, and so we predicted that predation would have a more severe effect on the reproductive success of dulled females.

**Figure 1.**
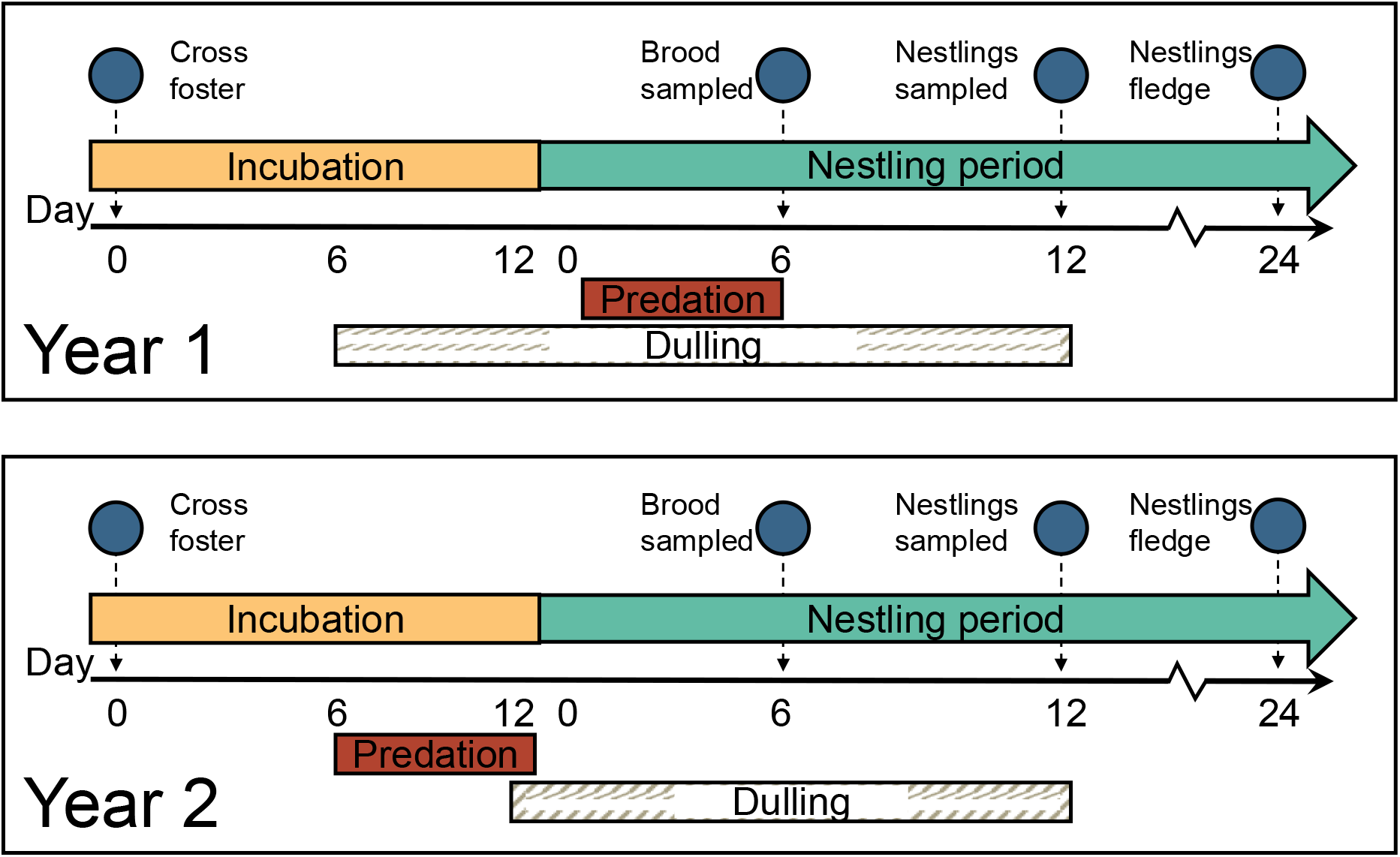
Schematic depicting the experimental design in year 1 (2018) and year 2 (2019) of the study.

## METHODS

We studied wild tree swallows breeding in nest boxes near Ithaca, New York, USA (42.503° N, 76.437° W) from May to July of 2018 and 2019. This population has been monitored continuously since 1986 using standardized field methods (Winkler et al., 2020). We conducted a separate experiment in each year; however, general methods for monitoring reproductive behavior were the same in each year except for where noted. Nest boxes were monitored every other day starting at the beginning of the breeding season and active nests were checked every day around the expected hatching date to determine the timing of clutch initiation, onset of incubation (+/− 1 day) and hatching (exact day, approximately 12 days after clutch completion).

### Experimental manipulations

In 2018 we carried out a 2×2 factorial experiment in which we first manipulated signal coloration of breeding female swallows and then later imposed a simulated predation challenge (Figure 1). At around day 6 of incubation, females were alternately assigned a “dulled” or control treatment. For this signal manipulation treatment, dulled females were colored across their entire white ventral surface with a light grey non-toxic marker (Faber-Castell PITT artist pen ‘big brush’ warm grey III 272). We previously validated that this treatment maintains the spectral characteristics of the plumage patch while reducing overall brightness (Taff et al., 2021). As a control, we applied a colorless marker over the same plumage area for the same length of time (Prismacolor Premier Colorless Blender PB-121; Newell Brands, Oak Brook, IL, U.S.A.). The treatments were re-applied one day after hatching and again six days after hatching so that the signal manipulation lasted during most of their reproductive attempt. In 2018, 20 females were experimentally dulled, and 22 females received the control treatment. We balanced treatment within age groups (second year vs. after second year) because breeding phenology and reproductive success differ between these ages in tree swallows (Winkler et al., 2020).

The second part of the experiment simulated an attempted predation event. For nests in the “predation” treatment, we simulated attempted predation on the female swallow by a mink (*Neovision vision*), which is a common predator of both adults and nestlings at our field sites. Females were trapped in the nest box and then gently pulled out of the box using a taxidermied mink wrapped around the researcher’s hand. The bird was brought to the ground below the nest box and then allowed to escape. During this treatment, the researcher’s face and body were covered with a camouflage suit and the female was held facing away from the researcher’s body to make the predation experience seem as realistic as possible. The predation simulation was performed three times during days 2-5 after hatching. The control group received no additional treatment outside of the signal manipulation (dulling or control) described in the previous paragraph. As above, we alternately assigned the predation and control treatments, while balancing these treatments within age classes. In 2018, 22 females were in the predation group and 20 females were in the control group.

In 2019 we repeated these experiments, but we reversed the order of the signal manipulation and predation treatments. Females were assigned to either predation or control treatments at day 6 of incubation and females in the predation treatment received two additional simulated predation attempts between days 8 and 12 of incubation. We included 29 females in the predation group and 33 in the control group. Then, on day 12-13 of incubation females were alternately assigned to either a plumage dulling or control signal manipulation treatment. Signal manipulation treatments were applied exactly as described for 2018, with coloring re-applied at the third capture on day 6 after hatching. In 2019, 31 females were experimentally dulled, and 31 females received the control treatment (Table S1).

### Nestling cross fostering and measurements

Differences in initial female quality are known to have a large effect on reproductive performance in tree swallows (Winkler et al., 2020). While our randomly assigned experimental treatments should account for these differences, we also sought to separate the effects of our treatments from any pre-treatment maternal effects by cross fostering eggs at each nest in the study. Nests were paired by breeding stage and on the fourth day of the egg laying stage we swapped half of the eggs from each nest and marked the bottom of all eggs with a pencil. For half of the nest pairs, we swapped an additional unmarked egg on the next day. This scheme ensured that egg-laying order was not associated with cross fostering status. In a few cases, we modified the swapping scheme to include three nests when appropriately timed matches were not available and a few later season nests were not cross fostered. For all nests, the ultimate clutch size remained the same after cross fostering.

Nestling growth and physiology at each nest were monitored as follows: On day 6 after hatching, we took a ‘total brood mass’ with all nestlings counted and weighed together (nearest 0.5 g). This total brood mass was divided by the number of nestlings to calculate a mean nestling mass at day 6. On day 12 after hatching, we banded nestlings with a USGS aluminum band and measured each nestling individually. We measured head + bill length (to the nearest 0.1 mm), flat wing length (to the nearest 0.5 mm), and mass (to the nearest 0.25 g). All blood samples were collected by brachial venipuncture into a heparinized micro-hematocrit tube. A baseline sample (< 70 μl) was collected within 3 minutes of capture followed by a stress-induced sample (< 30 μl) collected after 30 minutes of restraint. Immediately after the stress-induced sample was taken, we injected birds with 4.5 μl g^-1^ of dexamethasone to stimulate negative feedback (Mylan^®^ 4mg ml^-1^ dexamethasone sodium phosphate, product no.: NDC 67457-422-00).

All blood samples were stored on ice in the field for < 3 hours and then red blood cells and plasma were separated by centrifugation. Red blood cells were divided: part of the sample was stored in Longmire lysis buffer at room temperature for genotyping (Longmire et al., 1997). The other part was stored in NBS buffer (90% newborn calf serum and 10% DMSO) for telomere analysis. Samples for telomeres were kept at −80°C until analysis. Plasma was stored at −30°C until processing. We measured corticosterone with enzyme immunoassay kits (DetectX Corticosterone, Arbor Assays: K014-H5) that were previously validated for tree swallows in this population (see supplementary methods for details on extractions and hormone measurements; Taff et al., 2019a).

We used blood samples from nestlings to determine the nest of origin using a previously validated set of 9 microsatellite markers (Hallinger et al., 2019; Makarewich et al., 2009). For the purposes of this study, we were only interested in assigning nestlings to their correct mother from 2-3 possible females. Females were considered good matches if they matched nestlings at 8 of 9 loci and we were able to assign maternal origin for 374 of 386 nestlings sampled. Details on the lab procedure and criteria for assigning nest of origin can be found in the supplemental materials.

### Telomere quantification

We quantified relative nestling telomere length using a quantitative real-time PCR protocol following methods described in (Taff and Freeman-Gallant, 2017) and using current best practices (Morinha et al., 2020). Briefly, we extracted DNA from erythrocytes preserved in NBS buffer using Qiagen DNeasy Blood and Tissue Kits (Catalogue #69504, Valencia, CA). We used a QuickDrop spectrophotometer (Molecular Devices, San Jose, CA) to assess the DNA concentration and purity. The mean A260/280 absorbance ratio was 1.96 and the mean A260/230 absorbance ratio was 1.58. Because the A260/230 ratios tended to be lower than the mean recommended value (~1.8; Morinha et al., 2020) we tested for a correlation between sample A260/230 and the T/S ratio. We found no indication that the absorbance ratio was influencing our estimate of telomere length. We verified DNA integrity by running a subset of samples (~25%) on a 2% agarose gel; in all cases the DNA formed a single bold band with high molecular mass.

qPCR reactions were run on 384 well plates in reaction volumes of 13.5 μl. Each reaction contained 7 μl of PerfeCTa^®^ SYBR^®^ Green SuperMix, Low ROX™ (Quantabio, Beverly, MA), 2.8 picomoles of each primer and 14 ng of sample DNA. We amplified telomeres using qPCR with the primers Tel1b (5’- CGGTTTGTTTGGGTTTGGGTTTGGGTTTGGGTTTGGGTT-3’) and Tel2b (5’-GGCTTGCCTTACCCTTACCCTTACCCTTACCCTTACCCT-3’) which have been optimized for birds (Criscuolo et al., 2009) We amplified a single copy control gene (GAPDH: glyceraldehyde-3-phosphate dehydrogenase) using the primers GAPDH-F (5’-TTGACCACTGTCCATGCCATCAC-3’) and GAPDH-R (5’-TCCAGACGGCAGGTCAGGTC-3’). Both GAPDH and telomere reactions were run on a ViiA 7 Real-Time PCR System (Thermo Fisher Scientific, Waltham, MA). The telomere thermocycling conditions were as follows: 95°C for 10 min then 28 amplification cycles (95°C for 15s, 58°C for 30s, 72°C for 30s), followed by a melt curve (95°C for 15s, 60°C for 60s, 95°C for 15s). The GAPDH thermocycling conditions were as follows: 95°C for 10s, then 40 amplification cycles (95°C for 30s, 60°C for 30s), followed by a melt curve (95°C for 15s, 60°C for 60s, 95°C for 15s). Samples were run in triplicate for each reaction (telomere or GAPDH). Each plate also included three negative controls, a calibrator or “golden” sample (run on each plate to control for inter-plate variation) and five serial dilutions of a single high-quality sample. The repeatability of our standards across plates was 0.93 for both the telomere and GAPDH reactions.

### Telomere data processing

We exported raw florescence data from the Thermo Fisher Scientific Design and Analysis Software (v. 2.4.3) and then used LinRegPCR (v. 1.5.3) (Untergasser et al., 2021) to analyze amplification curves and calculate per-well efficiency and the quantification cycle (C_q_; the cycle number at which florescence rises above the threshold). Per-well reaction efficiencies ranged from 1.76 to 1.86 for the telomere reactions and from 1.87 to 2.03 for the GAPDH reaction (an efficiency value of 2 indicates the amount of the amplicon doubled each cycle). We examined the C_q_ values of triplicates to ensure precision in our estimation of telomere length. We averaged C_q_ values for each sample, including only replicates whose C_q_ values were within 0.25 standard deviations of one another. If we did not have at least two replicates within 0.25 standard deviations, we excluded or re-ran the sample.

We calculated the relative telomere length (RTL) for each sample using the following equation:

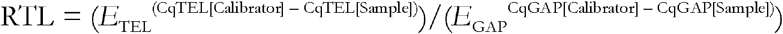

Where *E* is the mean reaction efficiency across all samples on a given plate; Cq[Calibrator] is the mean Cq across the calibrator samples on the plate, and Cq[Sample] is the mean Cq of a given sample (Reichert et al., 2017).

### Analysis

We used linear mixed models (LMMs) and generalized linear mixed models (GLMMs) to test for effects of the coloration and predation treatments on nestling growth and physiological measurements, including mass, skeletal measurements, corticosterone, and relative telomere length. Each response variable was modeled separately as a function of the fixed effects of color treatment, predation treatment, and their interaction. When the interaction term was not significant (all models), we removed it and report the results of the additive model. Models also included the covariate of female brightness before manipulation (numeric, centered and scaled). We initially included female brightness because this trait mediated the effect of plumage dulling in a previous experiment (Taff et al., 2021). We tested first for an interaction between initial brightness and color treatment; when the interaction effect was not significant, we removed the interaction and when the main effect of brightness was not significant, we removed the effect of brightness as well. We also initially included a covariate of whether the nestling was raised in its natal nest or was cross fostered (binary), but this effect was never significant, so we did not include it in any final model. Finally, models included two random effects: social nest box (i.e., the nest where the nestling was raised) and genetic mother. In some models the random effect of genetic mother did not explain any residual variance and caused a singular fit warning, indicating that the random effects structure was overfit.

In those cases, we removed the random effect of genetic mother, leaving only the random effect of social nest box. In R notation the full model structure is as follows: response_variable ~ predator_treatment + color_treatment * female_brightness + raised_nest + (1|nest_id) + (1|genetic_mom). Because of differences in experimental design each year, we ran separate models for 2018 and 2019. To test for differences in reproductive success between years, we ran GLMMs predicting per-nest hatching success and per-nest fledging success with year as a fixed factor, and nest id as a random effect.

We used mixed effects Cox proportional hazards models to test for differences in nestling survival between treatment groups. We included the fixed effects of color treatment and predation treatment and nest as a random effect. We tested for differences in overall fledging success using a logistic regression predicting nestling fate (fledged or died) based on treatment, with nest as a random effect.

## RESULTS

### Nestling growth and physiology

In 2018, when perceived predation risk was manipulated during the nestling period, the predation treatment had a negative effect on some measures of nestling growth. At 12 days of age, nestling wing length was significantly shorter in the predation group compared to controls (β = −8.07, CI = −14.08 – −2.06, *P* = 0.009; Table S2, Fig. 2). Nestling head + bill length was also smaller on average for predation nestlings compared to controls; however, difference was not significant (β = −0.91, CI = −1.86 – 0.04, *P* = 0.060; Table S3). Nestling mass at day 6 was significantly lower in the predation treatment compared to controls (β: −2.31, CI = −3.79 – −0.83, *P* = 0.004; Table S4). However, by day 12 mass was not significantly different between predation nestlings and control nestlings (β = −1.75, CI = −4.23 – 0.74, *P* = 0.166; Table S5).

**Figure 2:**
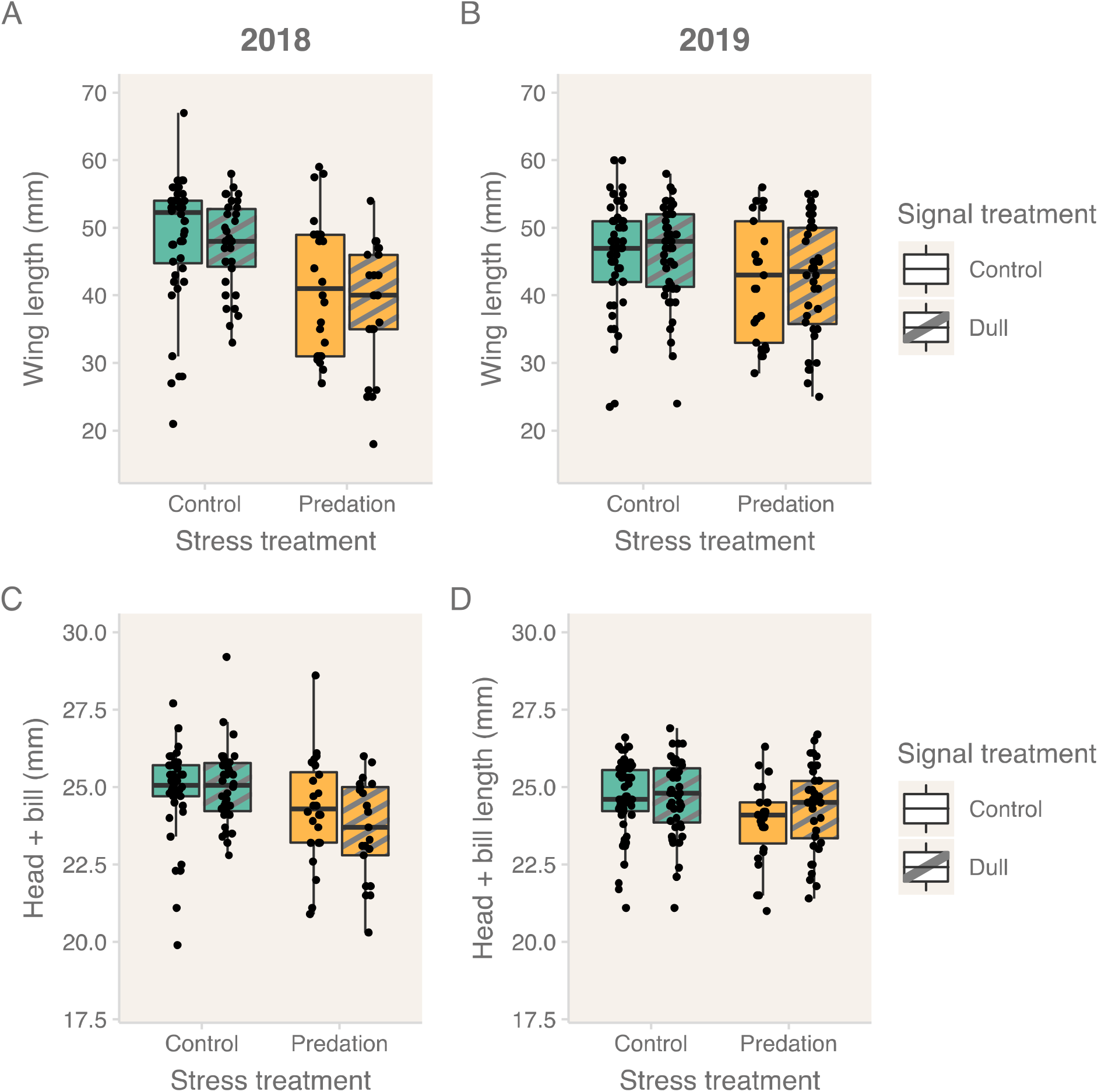
Nestling skeletal size at 12 days of age in predation and dulling treatments. Top row: wing length in A) 2018 and B) 2019. Bottom row: head + bill length in C) 2018 and D) 2019. Nestlings in the predation group (yellow) had significantly smaller wings at day 12 than nestlings in the control group (teal) in 2018. See results for full statistical comparisons.

In contrast, the dulling treatment (which occurred during incubation and the nestling period) in 2018 had only minor effects on nestling growth and effects depended on female brightness at the start of the season. There was no significant effect of dulling treatment on 12-day old nestling wing length, head + bill length, or mass (Fig. 2–3; Tables S2, S3, S5). There was a significant interaction between initial female brightness and the dulling treatment on six-day old nestling mass (Table S4): females in the experimentally dulled treatment had a positive relationship between pre-treatment brightness and nestling mass, whereas for females in the control treatment there was not a significant relationship between brightness and nestling mass (Fig. S1).

**Figure 3.**
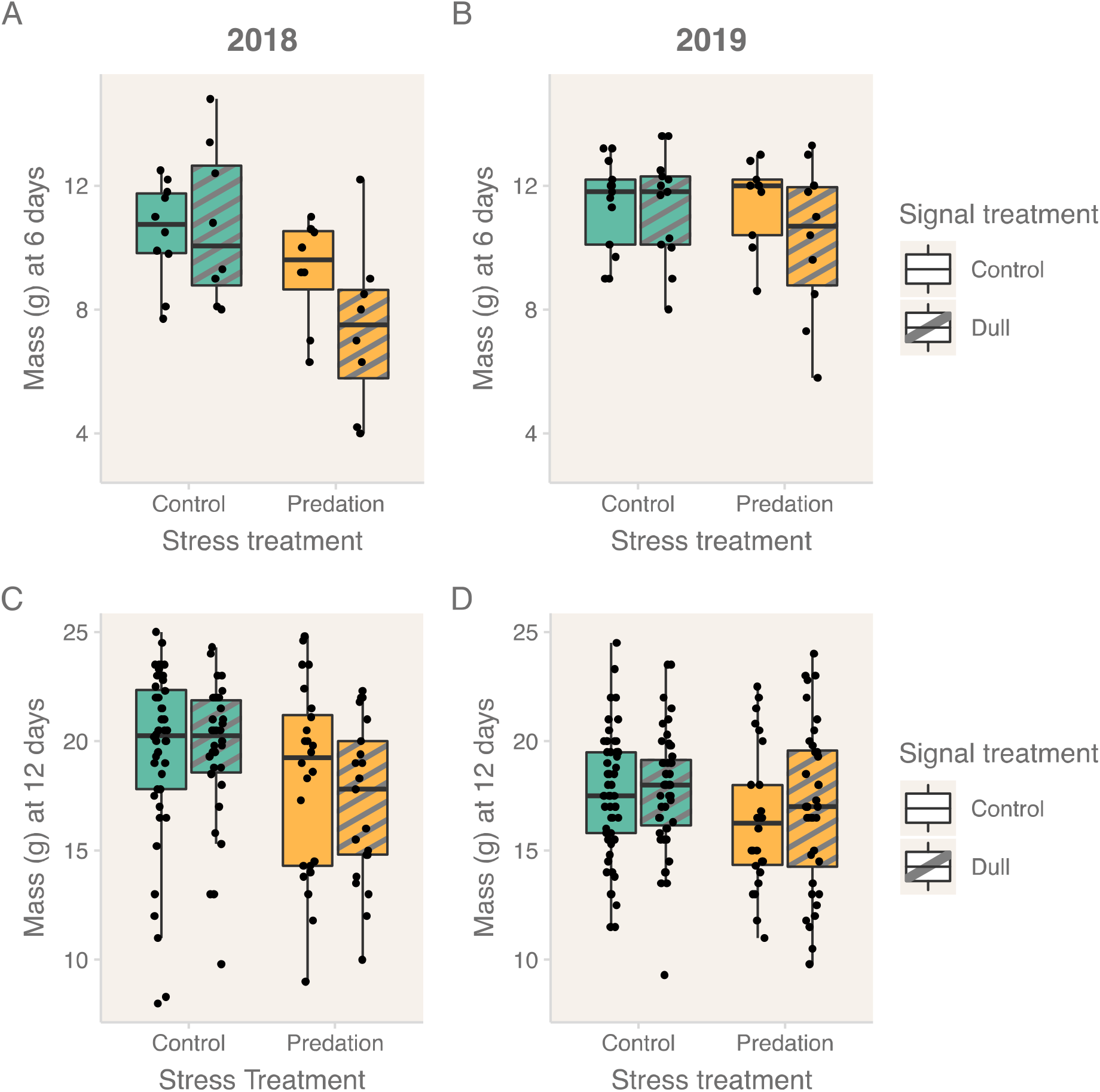
A) Average mass at 6 days of age in 2018 and B) 2019. C) Nestling mass at 12 days in 2018 and D) 2019. At 6 days of age all nestlings in the brood were massed together and then an average mass was calculated by dividing the total by the number of nestlings. At 12 days of age each nestling was individually massed. Each point in A-B is an average for a nest, each point in C-D is an individual nestling. Nestlings in the predation treatment (yellow) were significantly smaller than controls (teal) at 6 days of age in 2018. No other differences between treatment groups were significant.

In 2019, the predation treatment (which occurred during incubation) did not have a significant effect on nestling mass, wing length, or head + bill length (Fig. 2–3, Tables S2-5). However, the dulling treatment (which occurred during the nestling period) did have effects on nestling size, which were mediated by initial female brightness. There was a significant interaction between initial female brightness and the dulling treatment on nestling wing length (β = 4.93, CI = 0.31 – 9.54, *P* = 0.036), head + bill length (β = 0.73, CI = 0.09 – 1.37, P = 0.025) and 12-day nestling mass (β = 2.31, CI = 0.62 – 3.99, *P* = 0.008). For experimentally dulled females, nestling size was slightly positively correlated with initial female brightness. However, for control females, nestling size was negatively correlated with female brightness (Fig S1-2).

We tested for effects of the predation and dulling treatments on nestling corticosterone. Each year we quantified baseline, stress-induced, and post-dexamethasone corticosterone in 12-day old nestlings. There were no significant differences either between predator treatments or between dulling treatments in any corticosterone measurement (Fig. 4, Table S6-S7). In contrast to models of nestling size, there was no effect of initial female brightness on nestling corticosterone in either year.

**Figure 4.**
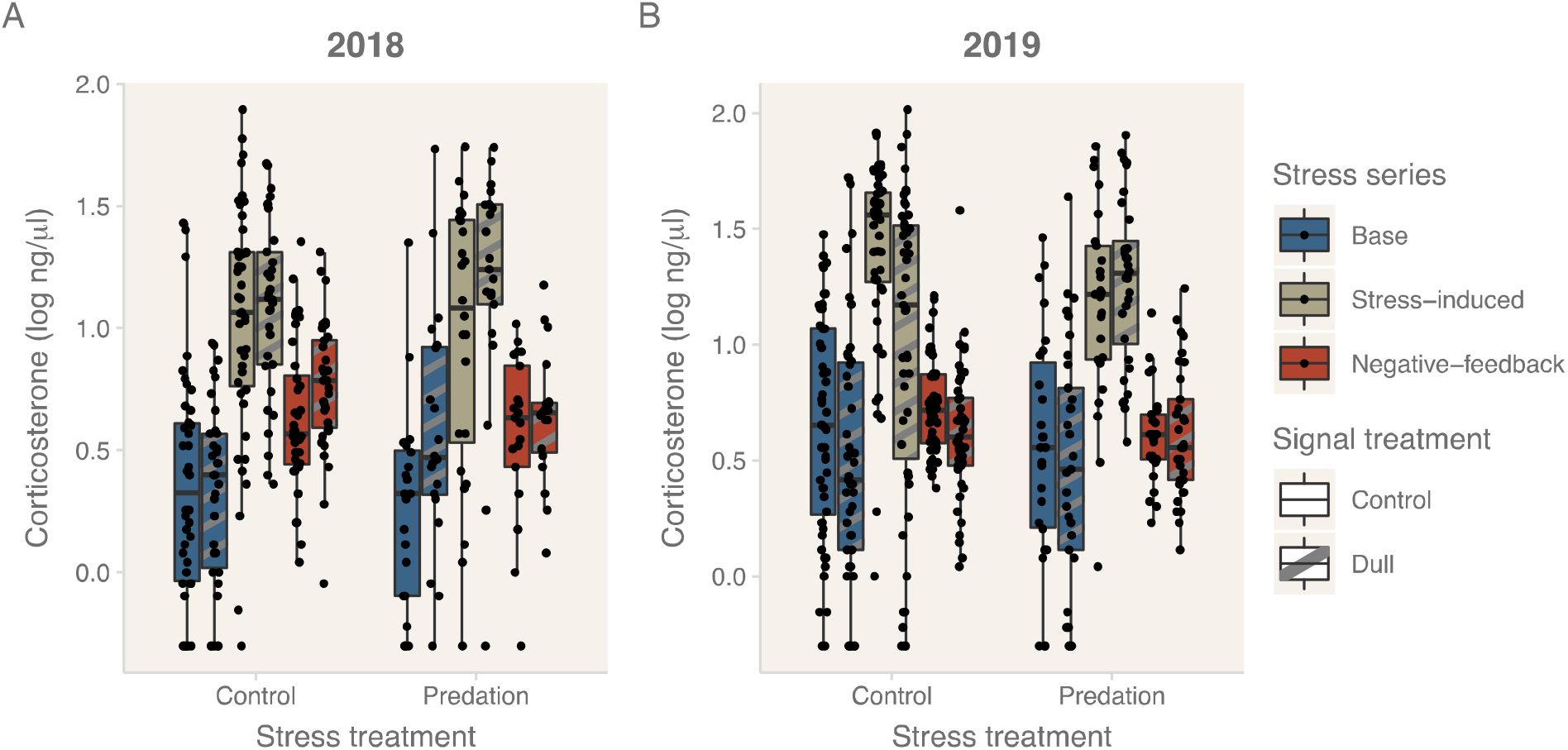
Corticosterone (cort) concentrations for nestlings at 12 days of age. We conducted a stress series for each nestling quantifying baseline (within three minutes of disturbance), stress-induced (after 30 minutes) and negative-feedback cort levels (30 minutes after injection with dexamethasone, see methods). There was no significant effect of either predation or signal manipulation in either year.

### Telomere length

Nestlings in 2018 had significantly shorter relative telomere lengths in the predation group compared to the control group (β = −0.05, CI = −0.10 – −0.01, *P* = 0.013; Fig. 5, Table S8). There was no significant difference between dulling and control treatments (β = 0.02, CI = −0.02 – 0.07, *P* = 0.270). In contrast, nestlings in 2019 had no significant difference in telomere lengths between predation and control groups (β = 0.05, CI = −0.01 – 0.12, *P* = 0.117). Again, there was no difference between dulling and control treatments (β = 0.00, CI =−0.06 – 0.06, *P* = 0.988; Fig. 5, Table S7). We tested whether relative telomere length was correlated with measurements of nestling quality. There was no relationship between telomere length and mass or wing length of 12-day old nestlings in either study year (Tables S9-S10). Telomere length also was not a significant predictor of fledging success in either year (Table S11); however, this analysis only included those nestlings that survived to 12 days of age (the date of blood sampling).

**Figure 5:**
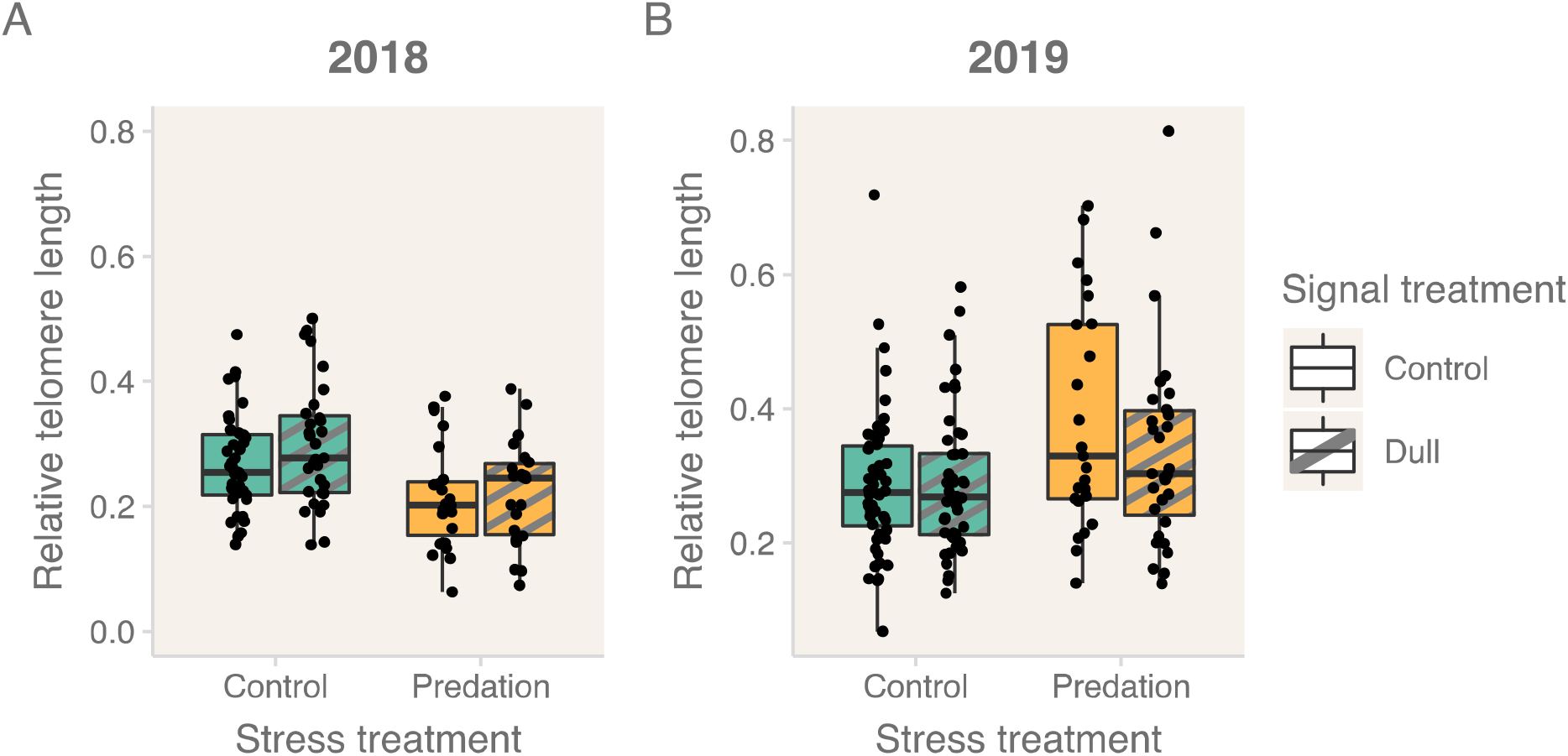
Relative telomere length in 2018 (A) and 2019 (B). Telomeres were significantly shorter in the predation group (yellow) compared to controls (teal) in 2018. There was no significant difference between predation and control groups in 2019 or between signal treatment groups in either year.

### Reproductive success

There was no effect of either the predation treatment or the dulling treatment on hatching success in either year (Table S12). We compared reproductive success (hatching success and fledging success) between our two study years to investigate whether environmental differences between years could have contributed to our results. Hatching success did not differ between years. In 2018, 81% of eggs hatched and in 2019, 87% of eggs hatched (Odds ratio = 1.55, CI = 0.55 – 4.39, *P* = 0.407). Once nestlings hatched, fledging success did not differ overall between years. In 2018, 51% of nestlings fledged and in 2019 52% of nestlings fledged (Odds ratio = 0.94, CI = 0.24 – 3.66, *P* = 0.932).

We used a Cox proportional hazards model to test for differences between treatments in nestling survival from hatching (day 0) to fledging (~ day 23). In 2018, nestlings in the predator treatment had a higher risk of death (hazard ratio = 3.38, CI = 1.32 – 8.62, *P* = 0.011, Fig. 6). In 2019, however, survival did not differ between predator and control treatments (hazard ratio = 2.07, CI = 0.72 – 5.96, *P* = 0.178, Fig. 6). There was no significant difference in either year between dulling treatments (Table S13). Fledging success was lower for nestlings in the predator treatment compared to the control treatment in 2018; however, it was not significantly different between predator and control treatments in 2019 (Table S14). There was no significant difference in fledging success between dulling and control treatments in either year (Table S14).

**Figure 6:**
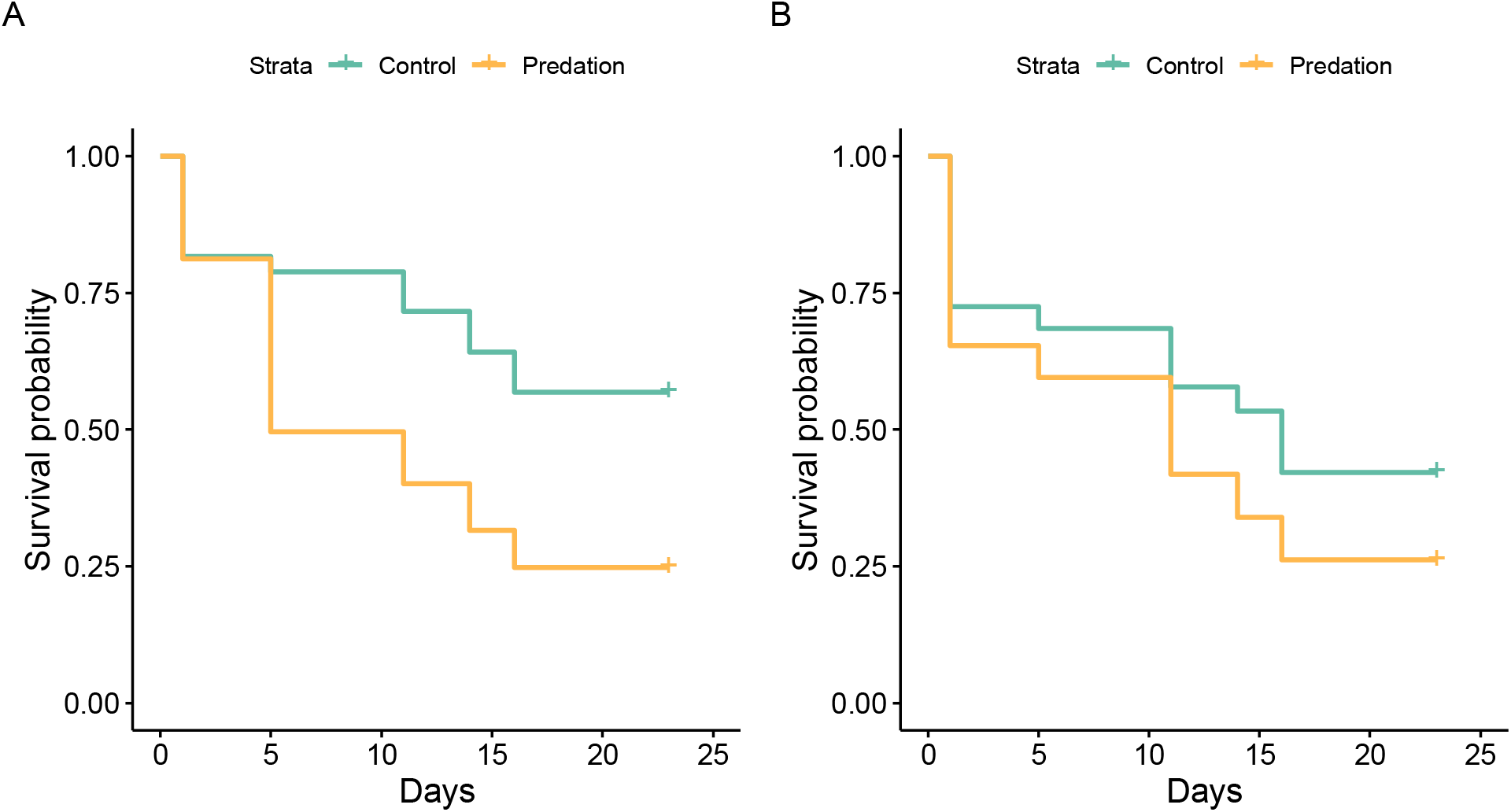
Daily survival probability of nestlings in the predator and control groups in 2018 (A) and 2019 (B). Shaded lines represent confidence intervals. Nests were checked at hatching (0 days), 6 days, 12 days, 15 days, and fledging (~23 days). In 2018 (A) simulated attempted predation occurred between days 1 – 5 of the nestling period. In 2018 (B) simulated attempted predation occurred before hatching (Fig. 1).

## DISCUSSION

The presence of predators affects bird physiology, behavior, and reproductive investment (Allen et al., 2022). Here, we tested for the effects of increased perceived predation risk on tree swallow reproductive success and sought to identify the physiological mechanisms involved in nestling stress resilience. We found that simulated predation attempts negatively affected tree swallows; however, the effects depended on the timing of predation events. During the first year of the study, when breeding females experienced three simulated predation attempts during the early postnatal period, their nestlings suffered lower fledging success and those that survived had reduced growth and shorter telomeres. In contrast, in the second year of the study, when predation attempts occurred during the prenatal period, there were no effects on nestling survival, growth, or physiology. Contrary to our predictions, signal manipulation (i.e., plumage dulling) did not mediate resilience to predation stress. The plumage dulling treatment did have minor effects on nestling growth, but only when it occurred during the nestling period and only after considering initial female brightness.

In 2018, the largest proportion (42%) of nestling deaths in the predation group occurred during the first six days of life, i.e., during or shortly after the experimental manipulation (Fig. 6). Tree swallow nestlings are unable to thermoregulate on their own until they are about 10 days old (Dunn, 1979); thus, reduced parental brooding to avoid predation may have caused nestling mortality. In addition to the immediate effects of predation stress on nestling survival, we observed secondary effects on nestling growth. At six days of age (shortly after predation events), nestlings in the predation group were significantly smaller in mass than those in the control group (Fig. 3). By 12 days of age there was no difference in mass between treatments, suggesting that nestlings may have been able to accelerate growth to recover from a temporary reduction in parental care. However, wing length at 12 days of age was still shorter in the predation group, indicating that not all aspects of development recovered. Previous studies that have manipulated the growth rate of nestlings have found that an acceleration in growth rate during development can have long term metabolic consequences for adults (Alonso-Alvarez et al., 2007; Criscuolo et al., 2008). Thus, even if swallows only temporarily reduce parental care in response to predators in the environment, the effects on growth and physiology of their offspring may be lasting.

In 2019, we reversed the order of the signal and predation manipulations and simulated predation before nestlings hatched. In contrast to the previous year, predation stress had no significant effect on nestling condition or survival. Nest abandonment in response to perceived predation risk can be high during incubation, when parental investment in reproduction is still relatively low (LaManna and Martin, 2016). Although hatching success was lower on average in the predation group, the differences were not statistically significant. Thus, our data indicate that breeding tree swallows are more resilient to simulated predation stress during the incubation period compared to the nestling period. Since eggs require less parental care than nestlings, behavioral changes of females during the incubation period are likely less consequential than similar changes during the first days of the nestling period.

We investigated the physiological mechanisms linking simulated predation to swallow fledging success. Direct contact with a predator can prompt a rise in nestling corticosterone (Herborn et al., 2014). The presence of predators may also indirectly provoke a hormonal response in nestlings through changes in parental care, e.g. reductions in provisioning or brooding (Crino et al., 2020; DuRant et al., 2010; Oers et al., 2015; Rensel et al., 2010). Although we expected that the effect would be most evident in nestlings, developing embryos may respond to predation stress as well. For example, yellow-legged gull eggs exposed to increased predator alarm calls hatched nestlings with elevated corticosterone levels and shorter telomeres (Noguera and Velando, 2019). Despite our predictions, we found no difference in corticosterone levels between predation and control groups in either year. There are several potential explanations for the lack of effect: First, the treatment may have only induced an acute response during the “predation” event or treatment period, and the response may have subsided by the time nestlings were sampled. Second, the HPA axis develops over the nestling period in altricial birds, and stressors early in development may not induce a detectible glucocorticoid response (Wada et al., 2009). Finally, corticosterone is just one of a number of hormones involved in the physiological response to a stressor, and corticosterone has physiological roles beyond the stress response (MacDougall-Shackleton et al., 2019). Indeed, the glucocorticoid response is not always predictable; other studies have similarly found no effect or unexpected effects of early life stressors on glucocorticoids in nestling birds (Ibáñez-Álamo et al., 2011; Wada et al., 2015). Correspondingly, there are increasing calls for a broader approach to characterizing the stress phenotype, including measures of oxidative stress, telomeres and other physiological traits (Bateson, 2016; MacDougall-Shackleton et al., 2019; Whitham et al., 2020).

In contrast to corticosterone, we found that simulated predation negatively affected telomere length of nestlings when predation events occurred early in the nestling period. In the first year of the study, nestlings in the predation treatment had shorter telomeres relative to controls (Fig. 5). Although telomere lengths were not associated with corticosterone concentrations, our results are consistent with previous studies showing that stressful conditions shorten telomeres, potentially through the negative effects of oxidative stress, and/or inflammation (Haussmann and Heidinger, 2015; Monaghan, 2014). Nestlings were not the direct targets of the simulated predation events; however, the stressor still had lasting negative effects on their skeletal size and telomere lengths. An earlier study at our field site found that first year telomere length in tree swallows predicts survival over the next three years (Haussmann et al., 2005). Thus, the indirect effects of predation stress on nestling tree swallows have the potential to affect long term survival, even for those birds that did fledge successfully.

We observed different effects of simulated predation stress in the two years of our study. While the most striking difference between our two field seasons was the timing of the experimental manipulations (pre- or post-hatching), it is possible that other factors may have contributed to the contrasting results. Tree swallow reproductive success is closely tied to environmental conditions and adverse weather events such as cold snaps reduce food availability and fledging success (Shipley et al., 2020). There was no significant difference in overall reproductive success between years, thus we do not expect that environmental differences between years were a primary driver of our results. Still, other factors, such as variation the density of birds in the area or abundance of natural predators could have contributed to the different results we observed each year.

The effects of the dulling treatment on tree swallow reproductive success were minor and did not affect the response to predation stress. Like the predation manipulation, plumage dulling had a stronger effect during the nestling period compared to the incubation period. In the second year of the study (2019), we found a significant effect of plumage dulling; however, the effect depended on initial female brightness. Earlier studies in our population and other populations of tree swallows found that females with brighter breast plumage are more resilient to environmental challenges and have higher reproductive success (Beck et al., 2015; Taff et al., 2019a). However, manipulation of this social signal sometimes leads to unexpected results. A previous study that experimentally dulled female plumage, as we did here, found that dulled females invested more in reproduction and had higher reproductive success (Taff et al., 2021). Manipulation of the breast plumage has the potential to create a “mismatch” between the social signal and the true quality of a female. Brighter females receive more aggressive interactions from conspecifics, and therefore may be forced to defend their territories more often (Coady and Dawson, 2013). In our study, we saw a positive relationship between initial brightness and nestling quality in the dulled females (Fig. S1). For females who were initially bright, dulling had a positive effect on the size of their nestlings (Fig. S3). It is possible that bright females experienced the advantages of bright plumage earlier in the breeding season while they were securing territories and mates but avoided negative conspecific interactions after they were dulled during the nestling period. On the other hand, for females that were initially duller than average, experimental dulling had a negative effect on the size of their nestlings at day 12 (Fig. S3). Experimental dulling of the lowest-quality females may have exacerbated any negative social position they had, further reducing their reproductive success.

Contrary to our initial predictions, plumage manipulation did not mediate resilience to predator stress. Interaction with the predator manipulation could have led to a “cancelling-out” effect of the dulling treatment. For instance, brighter females may have naturally been higher quality which increased their resilience to predation stress. However, experimentally dulled females may have been able to avoid aggressive interactions with other tree swallows, allowing them to invest more in reproduction and maintain fitness even under heightened predation stress. However, this theory does not explain why we saw no differences between dulling and control treatments in the absence of predation stress. Our sample sizes were slightly smaller than in Taff et al. 2021 (N = 34 – 36 per group in the previous study, N = 20 – 34 here) and so it is possible we did not have the power to detect small effects of the dulling treatment. Finally, it is also likely that the effects of the social environment are context-dependent and vary among years with environmental conditions, population density, and age/breeding condition of the females. The social environment of species like tree swallows likely interacts with HPA axis function and has the potential to mediate resilience to environmental stressors (Creel et al., 2013). However, isolating and manipulating these complex processes is difficult in wild populations and requires future investigation in our system.

## CONCLUSIONS

We found that the perceived risk of predation alters tree swallow reproductive success. Simulated predation events early in the nestling period resulted in increased nestling mortality, reduced size, and shorter telomeres. The effects of heightened perceived predation risk on nestling telomeres are especially notable because telomere length is linked to overall lifespan. Thus, our results add to a growing body of evidence demonstrating that transient, early-life stressors may have lasting effects, and that telomeres may link early-life conditions to later health and survival. Although previous studies have shown that the social environment may mediate how animals respond to stressors, here we did not find an interaction between our manipulation of a key social signal and the response of swallows to simulated predation. Birds live in dynamic environments involving challenges from intra- and inter-specific interactions. More work is needed in avian systems to understand the role of the social environment in a complex and changing world.

## ETHICAL NOTE

All procedures were approved by the Cornell University Institutional Animal Care & Use Board (IACUC protocol 2019-0023 and 2001-0051). Work was conducted under federal and state scientific collecting permits to MNV (USGS 24129, USFWS MB42428C; New York State 215 and 2350).

## DATA AVAILABILITY

Raw data and code for the analyses are available at https://github.com/smcnew/mcnew_etal_tres_nestling_telos and will be archived permanently on Zenodo upon acceptance.

## ACKNOWLEDGEMENTS

We thank the many field and lab assistants who helped with data collection including Bashir Ali, Allison Anker, Paige Becker, Raquel Castromonte, Jeremy Collison, KaiXin Chen, Alex Dopkin, Zapporah Ellis, Audrey Fox, Brianna Johnson, Christine Kallenberg, Raisa Kochmaruk, Alex Lee-Papastravos, Jabril Mohammed, Yusol Park, Alyssa Rodriguez, Bella Somoza, and Kwame Tannis. We also thank Brittany Laslow and Benj Sterrett for support in the lab and field. A.S.I, C.C.T., and S.M.M were funded by Edward Rose postdoctoral fellowships from the Cornell Lab of Ornithology. Additional funding for this research was provided by the U.S. Department of Defense, Defense Advanced Research Projects Agency (DARPA D17AP00033), the U.S. National Science Foundation (NSF IOS 1457251) and the U.S. National Institute of Food and Agriculture (NIFA Hatch 1017321) to M.V. The views, opinions and/ or findings expressed are those of the authors and should not be interpreted as representing the official views or policies of the U.S. Department of Defense or the U.S. Government

## Notes

https://github.com/smcnew/mcnew_etal_tres_nestling_telos

### Competing Interest Statement

The authors have declared no competing interest.

