## Supplementary materials for "Developmental stage-dependent effects of perceived predation risk on physiology and fledging success of tree swallows (*Tachycineta bicolor*)"

McNew et al 2022

**Additional details on methods**

***Determining Nest of Origin for Cross-Fostered Nestlings***

Blood samples collected in the field were stored in Longmire’s lysis buffer [1]. DNA was extracted from nestling and putative mother blood samples using Qiagen DNeasy Blood and Tissue kit spin columns following the manufacturer’s protocol. For each sample, we amplified a set of 9 microsatellite markers that were previously developed for and validated in this population of tree swallows [2]. These nine loci were amplified in two multiplexed reactions that included 5 and 4 loci each, as described in Hallinger et al. [3]. The full list of primer sequences, reaction volumes, and cycling conditions are available in Taff et al. [4]. After reactions were complete, we added 1 μl of the PCR product to 11.9 μl of formamide and 0.1 μl of LIZ size standard (GeneScanT M 500 LIZT M ) and submitted the samples to the Cornell Biotechnology Resources Center for fragment analysis. We re-ran any samples that failed to amplify and all 9 loci were scored for all nestlings and putative mothers in this study except for those noted below where a sample was not available. Peaks for each loci were called using the Microsatellite Plugin in Geneious version 11.0.4 with manual confirmation for each call. Because conspecific brood parasitism is rare in tree swallows [5], we assumed that putative mothers for each nestling included only the 2-3 females associated with nest boxes that were involved in each cross-fostering group. Therefore, we compared the peak calls for each nestling to the set of 2-3 putative mothers and identified matches. We considered nestlings to match a putative mother if their alleles matched at a minimum of 8 out of 9 loci and did not match the other putative mothers at a minimum of 2 out of 9 loci. Across the two years, 374 of 386 nestlings matched one female according to these criteria.

***Plumage Measurement***

We measured the overall brightness of feathers collected from the center of the white breast of each female in this study exactly as described in previous work on this population [6,7]. Briefly, four feathers from the breast were stacked and taped onto black construction paper. Reflectance was then measured with an Ocean Optics FLAME-S-UV-VIS spectrophotometer with PX-2 pulsed Xenon light source and WS-1 white standard in OceanView version 1.5.2 (Ocean Optics, Dunedin, FL). Acquisition settings included a 20 nm boxcar width, 10 scan average, and 60 ms integration time. We used a holster on the fiber optic UV/VIS probe so that light was blocked while reading and the probe was held a constant 5 mm distance from the feathers. For each feather stack, we took four separate spectral readings with the probe removed between each reading. We processed raw reflectance spectra using the pavo package in R [8]. As in Taff et al. [6], we focused on the overall brightness of the breast plumage given by the ‘B2’ measurement in pavo. This measurement represents the average reflectance across the range of 300-700 nm. Finally, the brightness measurements from the four separate spectra were averaged for each individual to arrive at a single brightness measure for each individual in the population before treatments were applied. In this study, we did not collect additional feathers at later time points after treatments, but Taff et al. [6]includes extensive validation data with feathers measured at varying time points after treatments for sham and

dulled birds.

***Corticosterone Measurement***

We measured the concentration of corticosterone from plasma that was frozen in the field using commercially available enzyme immunoassay (EIA) kits (DetectX Corticosterone, Arbor Assays: K014-H5). Validation and lab testing data on these kits when applied to our population of tree swallows is available in Taff et al. [6]. We first extracted corticosterone from plasma by adding 5 μl of plasma to 45 μl of assay buffer and then proceeding with three rounds of ethyl acetate extraction. The final extract was dried overnight in a fume hood and then reconstituted with 125 μl of assay buffer. Reconstituted samples were run in duplicate with a 9 point standard curve in 96 well EIA plates. Average extraction efficiency was determined by spiking some samples with a known amount of concentrated corticosterone and determining the percent recovery.

23

Using these starting volumes, the lower detection limit for corticosterone was 0.8 ng/μl and we substitute this value for any samples that were too low to detect. Overall, the extraction efficiency was 96.2%. When comparing replicates of each sample run on the same plate, the intra-plate CV was 11.5%. The inter-plate CV, based on plasma pools run across multiple plates, was 12.7%.

**Supplementary Tables**

Table S1: Sample sizes for experimental treatments in each year of the study.

|  |  | Simulated Predation | Control |
| --- | --- | --- | --- |
| Year 1 (2018) | |  |  |
|  | Plumage Dulling | 11 | 9 |
|  | Control | 11 | 11 |
| Year 2 (2019) | |  |  |
|  | Plumage Dulling | 15 | 16 |
|  | Control | 14 | 17 |

Table S2: Effects of treatment on nestling wing length in the two study years.

|  | **Wing length (2018)** | | | **Wing length (2019)** | | |
| --- | --- | --- | --- | --- | --- | --- |
| *Predictors* | *Estimates* | *CI* | *p* | *Estimates* | *CI* | *p* |
| Intercept (Controls) | 48.84 | 44.19 – 53.48 | **<0.001** | 45.48 | 42.07 – 48.90 | **<0.001** |
| Predation | -8.07 | -14.08 – -2.06 | **0.009** | -3.22 | -7.88 – 1.43 | 0.173 |
| Dull | -3.10 | -9.01 – 2.81 | 0.302 | 0.17 | -4.18 – 4.52 | 0.938 |
| Brightness (scaled) | 3.05 | -0.01 – 6.10 | 0.050 | -2.98 | -6.74 – 0.79 | 0.120 |
| Dull x Brightness |  |  |  | 4.93 | 0.31 – 9.54 | **0.036** |
| **Random Effects** | | | | | | |
| σ^2^ | 24.34 | | | 24.24 | | |
| τ_00_ | 4.65 _gen_mom_ | | | 7.04 _gen_mom_ | | |
|  | 56.91 _site_nest_ | | | 35.32 _site_nest_ | | |
| ICC | 0.72 | | | 0.64 | | |
| N | 31 _site_nest_ | | | 38 _site_nest_ | | |
|  | 39 _gen_mom_ | | | 51 _gen_mom_ | | |
| Observations | 121 | | | 161 | | |
| Marginal R^2^ / Conditional R^2^ | 0.243 / 0.786 | | | 0.101 / 0.673 | | |

Table S3. Effects of treatment on nestling head + bill length in the two study years.

|  | **Head-bill (2018)** | | | **Head-bill (2019)** | | |
| --- | --- | --- | --- | --- | --- | --- |
| *Predictors* | *Estimates* | *CI* | *p* | *Estimates* | *CI* | *p* |
| Intercept (Controls) | 24.93 | 24.21 – 25.66 | **<0.001** | 24.40 | 23.92 – 24.88 | **<0.001** |
| Predation | -0.91 | -1.86 – 0.04 | 0.060 | -0.43 | -1.06 – 0.20 | 0.179 |
| Dull | -0.24 | -1.17 – 0.69 | 0.613 | 0.30 | -0.27 – 0.88 | 0.300 |
| Brightness (scaled) | 0.53 | 0.05 – 1.01 | **0.031** | -0.62 | -1.14 – -0.11 | **0.018** |
| Dull x Brightness |  |  |  | 0.73 | 0.09 – 1.37 | **0.025** |
| **Random Effects** | | | | | | |
| σ^2^ | 0.87 | | | 0.75 | | |
| τ_00_ | 1.41 _site_nest_ | | | 0.38 _gen_mom_ | | |
|  |  | | | 0.48 _site_nest_ | | |
| ICC | 0.62 | | | 0.53 | | |
| N | 31 _site_nest_ | | | 38 _site_nest_ | | |
|  |  | | | 51 _gen_mom_ | | |
| Observations | 121 | | | 163 | | |
| Marginal R^2^ / Conditional R^2^ | 0.179 / 0.686 | | | 0.093 / 0.577 | | |

Table S4. Effect of treatment on average nestling mass per nest at 6 days of age*

|  | **Av. 6 day old mass (2018)** | | | **Av. 6 day old mass (2019)** | | |
| --- | --- | --- | --- | --- | --- | --- |
| *Predictors* | *Estimates* | *CI* | *p* | *Estimates* | *CI* | *p* |
| Intercept (Controls) | 10.87 | 9.67 – 12.07 | **<0.001** | 11.61 | 10.71 – 12.50 | **<0.001** |
| Predation | -2.31 | -3.79 – -0.83 | **0.004** | -0.52 | -1.61 – 0.58 | 0.347 |
| Dull | -0.69 | -2.18 – 0.79 | 0.347 | -0.52 | -1.61 – 0.56 | 0.337 |
| Brightness (scaled) | -0.07 | -1.05 – 0.90 | 0.876 |  |  |  |
| Dull x Brightness | 1.56 | 0.03 – 3.09 | **0.046** |  |  |  |
| Observations | 33 | | | 45 | | |
| R^2^ / R^2^ adjusted | 0.403 / 0.318 | | | 0.043 / -0.002 | | |

* At 6 days of age the entire brood was massed together. The average mass was calculated by dividing the total mass by the number of nestlings.

Table S5. Effect of treatment on nestling mass at 12 days of age*

|  | **2018** | | | **2019** | | |
| --- | --- | --- | --- | --- | --- | --- |
| *Predictors* | *Estimates* | *CI* | *p* | *Estimates* | *CI* | *p* |
| Intercept (Controls) | 19.79 | 17.85 – 21.74 | **<0.001** | 17.27 | 16.02 – 18.52 | **<0.001** |
| Predation | -1.75 | -4.23 – 0.74 | 0.166 | -0.80 | -2.51 – 0.91 | 0.358 |
| Dull | -0.66 | -3.11 – 1.79 | 0.595 | 0.62 | -1.00 – 2.24 | 0.454 |
| Brightness (scaled) |  |  |  | -1.51 | -2.90 – -0.12 | **0.034** |
| Dull x Brightness |  |  |  | 2.31 | 0.62 – 3.99 | **0.008** |
| **Random Effects** | | | | | | |
| σ^2^ | 5.97 | | | 4.65 | | |
| τ_00_ | 10.23 _site_nest_ | | | 4.97 _site_nest_ | | |
| ICC | 0.63 | | | 0.52 | | |
| N | 32 _site_nest_ | | | 38 _site_nest_ | | |
| Observations | 123 | | | 163 | | |
| Marginal R^2^ / Conditional R^2^ | 0.049 / 0.650 | | | 0.119 / 0.574 | | |

* At 12 days of age nestlings were massed individually.

Table S6. Models testing for effects of treatments on 12-day old nestling corticosterone in 2018

|  | **base cort** | | | **stress cort** | | | **post-dex cort** | | |
| --- | --- | --- | --- | --- | --- | --- | --- | --- | --- |
| *Predictors* | *Estimates* | *CI* | *p* | *Estimates* | *CI* | *p* | *Estimates* | *CI* | *p* |
| Intercept (Controls) | 2.71 | 0.46 – 4.96 | **0.019** | 16.84 | 9.79 – 23.90 | **<0.001** | 5.63 | 3.89 – 7.38 | **<0.001** |
| Predation | 1.77 | -1.17 – 4.71 | 0.236 | 2.47 | -6.55 – 11.49 | 0.588 | -1.19 | -3.48 – 1.09 | 0.302 |
| Dull | 1.26 | -1.62 – 4.14 | 0.388 | -0.70 | -9.60 – 8.21 | 0.877 | 1.02 | -1.21 – 3.26 | 0.366 |
| **Random Effects** | | | | | | | | | |
| σ^2^ | 34.26 | | | 100.74 | | | 10.66 | | |
| τ_00_ | 6.73 _site_nest_ | | | 127.13 _site_nest_ | | | 6.39 _site_nest_ | | |
| ICC | 0.16 | | | 0.56 | | | 0.37 | | |
| N | 32 _site_nest_ | | | 32 _site_nest_ | | | 31 _site_nest_ | | |
| Observations | 118 | | | 119 | | | 112 | | |
| Marginal R^2^/ Conditional R^2^ | 0.027 / 0.187 | | | 0.007 / 0.561 | | | 0.033 / 0.395 | | |

Table S7. Models testing for effects of treatments on 12-day old nestling corticosterone in 2019.

|  | **base cort** | | | **stress cort** | | | **post-dex cort** | | |
| --- | --- | --- | --- | --- | --- | --- | --- | --- | --- |
| *Predictors* | *Estimates* | *CI* | *p* | *Estimates* | *CI* | *p* | *Estimates* | *CI* | *p* |
| Intercept (Controls) | 7.44 | 3.68 – 11.19 | **<0.001** | 28.73 | 19.98 – 37.48 | **<0.001** | 6.27 | 3.35 – 9.18 | **<0.001** |
| Predation | -2.25 | -7.29 – 2.79 | 0.379 | -2.69 | -14.39 – 9.00 | 0.650 | -1.96 | -5.78 – 1.86 | 0.312 |
| Dull | -0.37 | -5.26 – 4.52 | 0.881 | -4.20 | -15.56 – 7.16 | 0.466 | 1.03 | -2.69 – 4.76 | 0.584 |
| **Random Effects** | | | | | | | | | |
| σ^2^ | 33.58 | | | 204.14 | | | 4.88 | | |
| τ_00_ | 46.90 _site_nest_ | | | 246.05 _site_nest_ | | | 30.26 _site_nest_ | | |
| ICC | 0.58 | | | 0.55 | | | 0.86 | | |
| N | 37 _site_nest_ | | | 37 _site_nest_ | | | 36 _site_nest_ | | |
| Observations | 155 | | | 156 | | | 150 | | |
| Marginal R^2^ / Conditional R^2^ | 0.015 / 0.589 | | | 0.014 / 0.553 | | | 0.031 / 0.865 | | |

Table S8. Effects of treatment on nestling telomere length

|  | **Relative telomere length (2018)** | | | **Relative telomere length (2019)** | | |
| --- | --- | --- | --- | --- | --- | --- |
| *Predictors* | *Estimates* | *CI* | *p* | *Estimates* | *CI* | *p* |
| Intercept (Controls) | 0.27 | 0.23 – 0.30 | **<0.001** | 0.29 | 0.24 – 0.35 | **<0.001** |
| Predation | -0.05 | -0.10 – -0.01 | **0.013** | 0.05 | -0.01 – 0.12 | 0.117 |
| Dull | 0.02 | -0.02 – 0.07 | 0.270 | 0.00 | -0.06 – 0.06 | 0.988 |
| **Random Effects** | | | | | | |
| σ^2^ | 0.00 | | | 0.01 | | |
| τ_00_ | 0.00 _gen_mom_ | | | 0.00 _gen_mom_ | | |
|  | 0.00 _site_nest_ | | | 0.01 _site_nest_ | | |
| ICC | 0.38 | | | 0.49 | | |
| N | 41 _gen_mom_ | | | 49 _gen_mom_ | | |
|  | 32 _site_nest_ | | | 37 _site_nest_ | | |
| Observations | 116 | | | 149 | | |
| Marginal R^2^ / Conditional R^2^ | 0.097 / 0.438 | | | 0.038 / 0.505 | | |

Table S9. Relationship between telomere length and nestling wing length.

|  | **Wing length 2018** | | | **Wing length 2019** | | |
| --- | --- | --- | --- | --- | --- | --- |
| *Predictors* | *Estimates* | *CI* | *p* | *Estimates* | *CI* | *p* |
| Intercept (Controls) | 45.72 | 40.60 – 50.83 | **<0.001** | 46.04 | 42.57 – 49.51 | **<0.001** |
| relative telomere length | -8.16 | -23.88 – 7.55 | 0.306 | -3.12 | -11.55 – 5.32 | 0.466 |
| **Random Effects** | | | | | | |
| σ^2^ | 32.52 | | | 24.77 | | |
| τ_00_ | 73.59 _site_nest_ | | | 40.06 _site_nest_ | | |
| ICC | 0.69 | | | 0.62 | | |
| N | 32 _site_nest_ | | | 37 _site_nest_ | | |
| Observations | 117 | | | 151 | | |
| Marginal R^2^ / Conditional R^2^ | 0.005 / 0.695 | | | 0.003 / 0.619 | | |

Table S10. Relationship between nestling telomere length and mass

|  | **Mass 2018** | | | **Mass 2019** | | |
| --- | --- | --- | --- | --- | --- | --- |
| *Predictors* | *Estimates* | *CI* | *p* | *Estimates* | *CI* | *p* |
| Intercept (Controls) | 17.59 | 16.22 – 18.95 | **<0.001** | 19.48 | 17.47 – 21.49 | **<0.001** |
| relative telomere length | -0.16 | -3.63 – 3.31 | 0.928 | -2.49 | -8.87 – 3.90 | 0.442 |
| **Random Effects** | | | | | | |
| σ^2^ | 4.38 | | | 5.49 | | |
| τ_00_ | 5.05 _site_nest_ | | | 9.91 _site_nest_ | | |
| ICC | 0.54 | | | 0.64 | | |
| N | 37 _site_nest_ | | | 32 _site_nest_ | | |
| Observations | 153 | | | 117 | | |
| Marginal R^2^ / Conditional R^2^ | 0.000 / 0.535 | | | 0.003 / 0.645 | | |

Table S11. Relationship between telomere length and fledging success

|  | **Fledging success 2018** | | | **Fledging success 2019** | | |
| --- | --- | --- | --- | --- | --- | --- |
| *Predictors* | *Odds Ratios* | *CI* | *p* | *Odds Ratios* | *CI* | *p* |
| Intercept (Controls) | 13.97 | 0.84 – 231.89 | 0.066 | 19.83 | 1.08 – 365.55 | **0.045** |
| relative telomere length | 0.10 | 0.00 – 331.61 | 0.574 | 0.06 | 0.00 – 28.56 | 0.364 |
| **Random Effects** | | | | | | |
| σ^2^ | 3.29 | | | 3.29 | | |
| τ_00_ | 7.50 _site_nest_ | | | 11.63 _site_nest_ | | |
| ICC | 0.70 | | | 0.78 | | |
| N | 31 _site_nest_ | | | 37 _site_nest_ | | |
| Observations | 112 | | | 153 | | |
| Marginal R^2^ / Conditional R^2^ | 0.004 / 0.696 | | | 0.009 / 0.782 | | |

Table S12. Differences in hatching success between treatments each year

|  | **Hatching success 2018** | | | **Hatching success 2019** | | |
| --- | --- | --- | --- | --- | --- | --- |
| *Predictors* | *Odds Ratios* | *CI* | *p* | *Odds Ratios* | *CI* | *p* |
| Intercept (Controls) | 34.80 | 2.61 – 463.30 | **0.007** | 10.68 | 4.59 – 24.85 | **<0.001** |
| Predation | 0.84 | 0.08 – 9.04 | 0.884 | 1.40 | 0.54 – 3.60 | 0.487 |
| Dulling | 0.86 | 0.08 – 9.29 | 0.900 | 0.56 | 0.22 – 1.42 | 0.224 |
| N | 42 _site_nest_ | | | 48 _site_nest_ | | |
| Observations | 42 | | | 48 | | |

Table S13. Mixed-effects Cox PH model predicting the effects of treatment on nestling survival.

|  | **Hazard ratio nestling survival 2018** | | | **Hazard ratio nestling survival 2019** | | |
| --- | --- | --- | --- | --- | --- | --- |
| *Predictors* | *Estimates* | *CI* | *p* | *Estimates* | *CI* | *P* |
| Predation | 3.38 | 1.32 – 8.62 | **0.011** | 2.07 | 0.72 – 5.96 | 0.178 |
| Dulling | 1.10 | 0.43 – 2.78 | 0.845 | 1.17 | 0.38 – 3.59 | 0.787 |
| N | 42 _site_nest_ | | | 62 _site_nest_ | | |
| Observations | 226 | | | 331 | | |

Table S14. Effects of treatment on fledging success in each year of the study

|  | **Fledging success 2018** | | | **Fledging success 2019** | | |
| --- | --- | --- | --- | --- | --- | --- |
| *Predictors* | *Odds Ratios* | *CI* | *p* | *Odds Ratios* | *CI* | *p* |
| Intercept (Controls) | 0.62 | 0.19 – 2.04 | 0.430 | 2.94 | 0.76 – 11.28 | 0.117 |
| Predation | 10.61 | 2.33 – 48.31 | **0.002** | 3.02 | 0.60 – 15.16 | 0.180 |
| Dulling | 0.86 | 0.21 – 3.54 | 0.836 | 1.01 | 0.21 – 4.83 | 0.987 |
| Brightness |  |  |  | 0.38 | 0.17 – 0.86 | **0.021** |
| **Random Effects** | | | | | | |
| σ^2^ | 3.29 | | | 3.29 | | |
| τ_00_ | 3.54 _site_nest_ | | | 6.59 _site_nest_ | | |
| ICC | 0.52 | | | 0.67 | | |
| N | 42 _site_nest_ | | | 62 _site_nest_ | | |
| Observations | 226 | | | 331 | | |
| Marginal R^2^ / Conditional R^2^ | 0.169 / 0.600 | | | 0.123 / 0.708 | | |

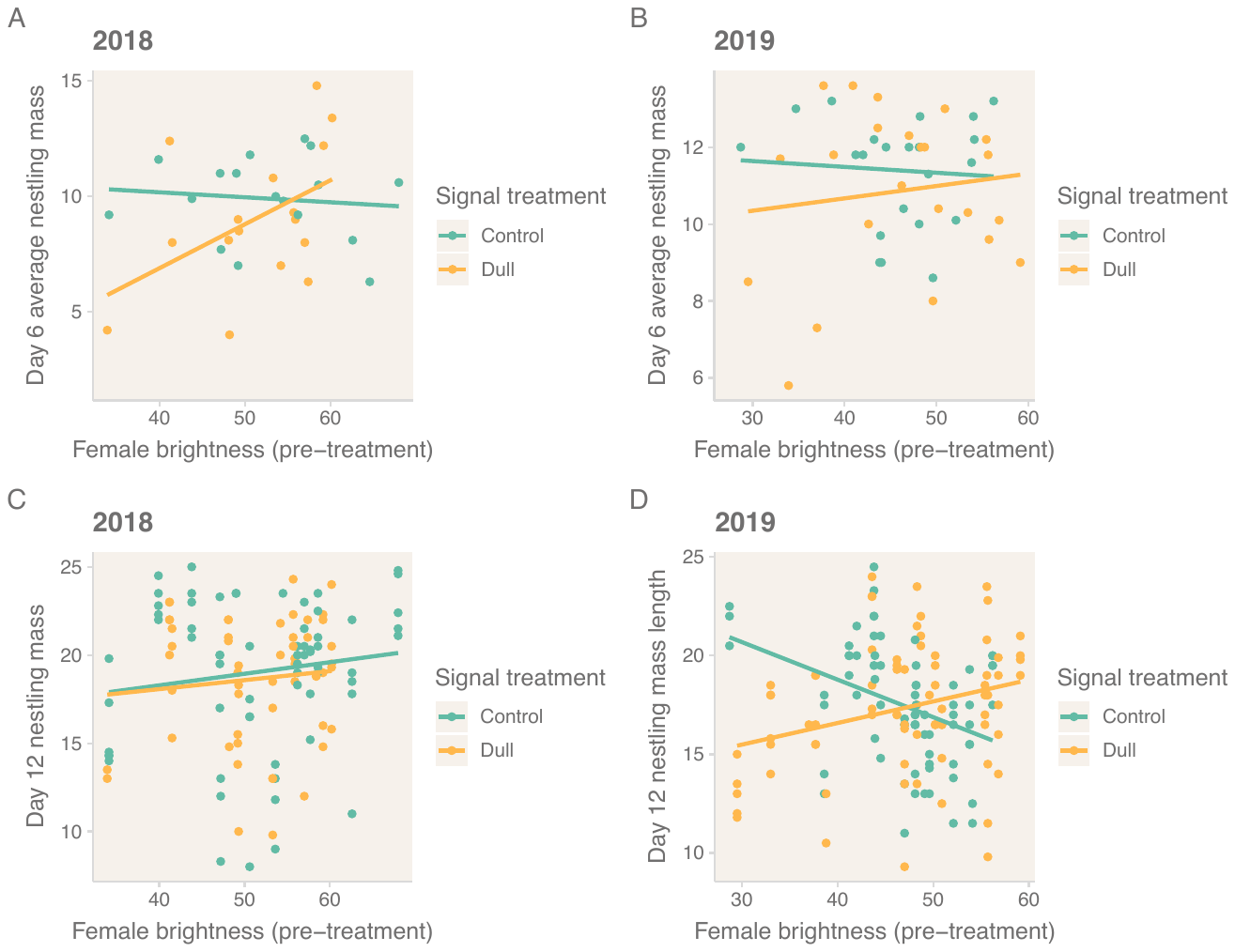

Figure S1. Relationship between female brightness before dulling treatment and mass of nestlings. In 2018, at six days of age there was a significant interaction between brightness and dulling treatment (P = 0.046; Table S3). In 2019, there was a significant interaction between brightness and dulling treatment at 12 days of age (P = 0.004; Table S4).

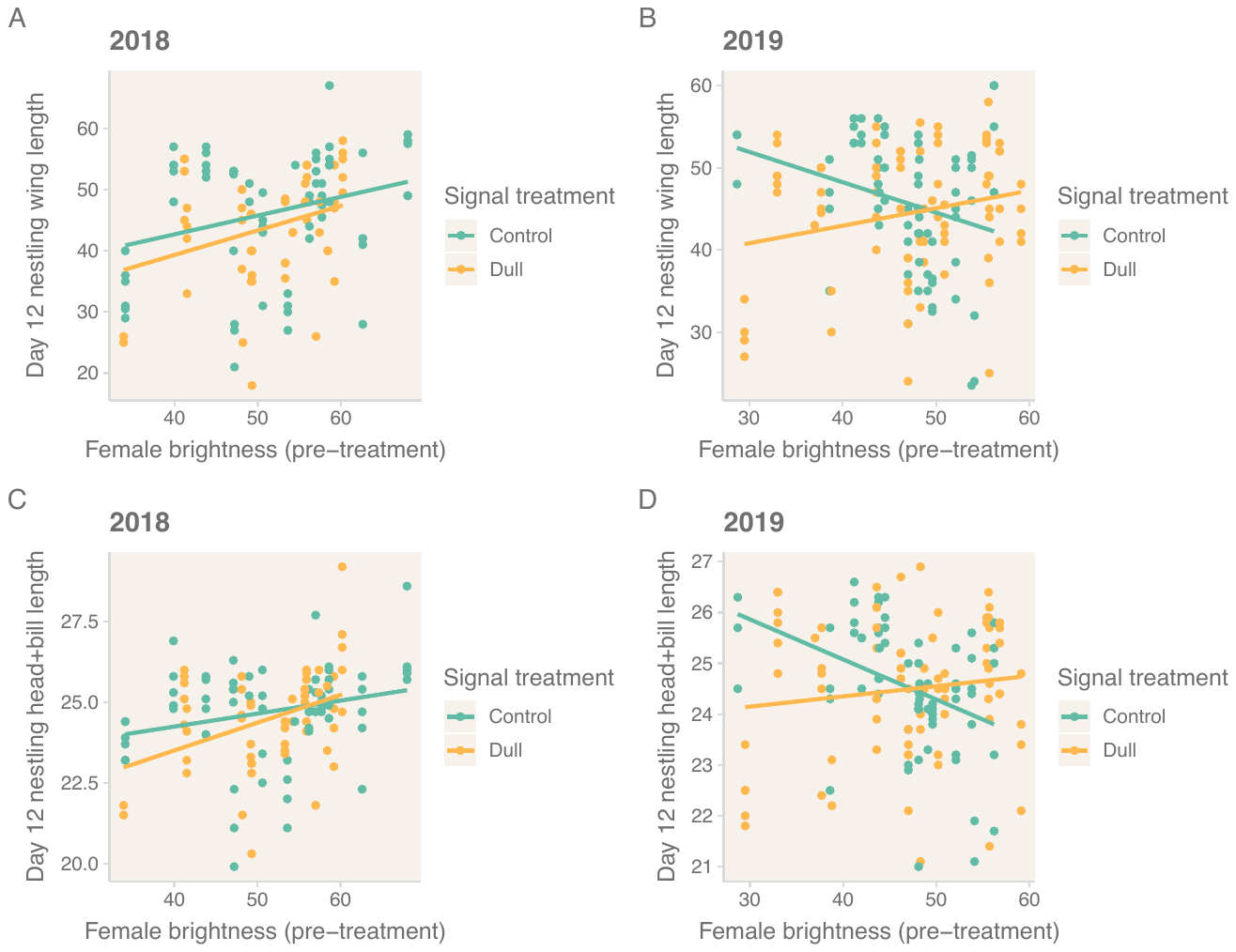

Figure S2. Relationship between female brightness before dulling treatment and skeletal size of nestlings. In 2018, brighter females had significantly larger nestlings, regardless of treatment group (control or dulled, Table S1-S2). In 2019, there were significant interactions between dulling treatment and brightness. For experimentally dulled females, nestling size was slightly positively correlated with initial female brightness. However, for control females, nestling size was negatively correlated with female brightness.
